# Cryo-EM structure of shutdown human non-muscle myosin 2A

**DOI:** 10.1101/2025.10.15.682586

**Authors:** David Casas-Mao, Glenn Carrington, Michelle Peckham

## Abstract

Determining the high-resolution structure of the widely expressed non-muscle myosin 2A (NM2A), in its dephosphorylated shutdown state, is important in understanding its regulation and disease roles. In shutdown molecules, the coiled-coil tail wraps around the myosin heads preventing them from forming filaments and binding to actin. We have solved the shutdown structure of NM2A to a global resolution of 3.0 Å in the heads region, and 6.3 Å for the whole molecule. This reveals specific ionic interactions that explain why the path of the coiled coil and the shutdown mechanism for NM2A differs from that of β-cardiac myosin and provides key insight into how specific mutations likely destabilize the shutdown state, leading to disease.

## Introduction

Myosins, molecular motors that move along F-actin filaments driven by ATP hydrolysis, are comprised of a complex of heavy chains and light chains. The N-terminal region of the heavy chain comprises the motor domain, which contains both nucleotide and F-actin binding sites. The motor is followed by the lever which, in all class 2 myosins, comprises the converter domain followed by an α-helix containing two IQ motifs, to which one essential (ELC) and one regulatory (RLC) light chain binds. The lever is followed by a long stretch of coiled coil in class 2 myosins, which both dimerizes the two heavy chains, and enables filament assembly, through its repeating patterns of charged residues along its length (*1*).

Smooth and non-muscle class 2 myosins switch between an active filamentous state and individual shutdown (10S) molecules (*2–4*). In the shutdown state, the coiled-coil tail wraps completely around the heads and the ATPase of these molecules is very low (*5*). Phosphorylation of the regulatory light chain (RLC), primarily at S19 by myosin light chain kinase, disrupts the shutdown state, allows myosin to extend and then assemble into filaments (*6–8*).

Relatively high-resolution structures for shutdown smooth muscle myosin (SMM) (up to 3.4Å) in the myosin heads region have been reported (*9–11*) and used to build a homology model for NM2A (*4*). However, a high-resolution structure for shutdown non-muscle myosin 2A (NM2A) has not yet been reported. NM2A is the only isoform of class 2 myosin found in platelets (*12*). On platelet activation, RLC phosphorylation rapidly increases from ∼10% (*13*) to about 90% within ∼1 minute (*14*), driving filament formation and platelet spreading, demonstrating the critical role of RLC phosphorylation in the biological function of NM2A. Critically, over ∼200 autosomal dominant missense mutations have been reported for the gene (MYH9) that encodes NM2A heavy chain, which cause bleeding disorders, kidney abnormalities, hearing defects and other disorders (*15–17*). Thus, we set out to solve the structure for shutdown NM2A to determine how it forms the shutdown state and to aid interpretation of disease mutations.

## Results

Using CryoEM, we solved the structure of dephosphorylated human NM2A to a global resolution of 3.0 Å in the heads region (Fig. 1A-D, Fig.S1A-G). The overall organization of the shutdown structure (Fig. 1A-D) is similar to that previously reported for SMM structures obtained by CryoEM (*9–11*). The structure of the motor domains reveals that both contain ADP.Pi (Fig. S2A-D) and that the backdoor, comprised of residues R234 and E457, is shut, preventing Pi release (Fig. S2E-F). The overall organization of the nucleotide binding pocket is similar to that recently reported for sequestered auto-inhibited interacting heads structure of β-cardiac myosin (β-CM) (*18*) (Fig. S2G) and is unlike that claimed for shutdown smooth muscle myosin (*9*).

**Fig. 1.**
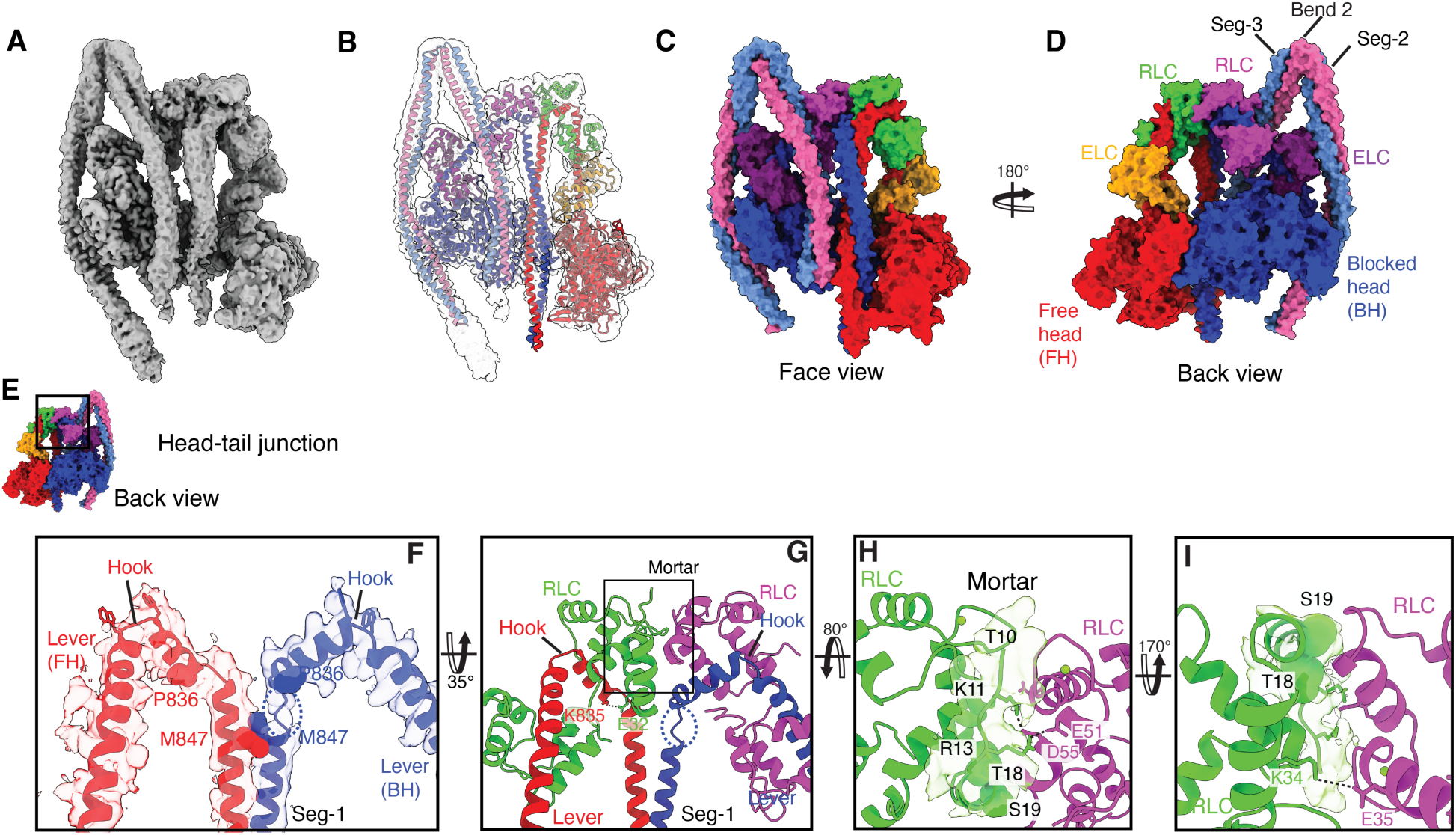
Structure of the interacting heads region of shutdown NM2A and contributions of the free head RLC to the shutdown state. (**A**) Cryo-electron density map of the heads region (Face view; contour level = 0.02) (**B)**. Structure of the heads region fit into the density map. Surface representations of the front (**C**) and back (**D**) views of the structure. The heavy chain for the blocked head (BH) is colored blue, and for the free head (FH) is colored red; the essential light chain (ELC) is colored dark purple, for the blocked head, and yellow for the free head. The regulatory light chain (RLC) is colored magenta for the BH and green for the FH. The boxed region in (**E)**, showing the head-tail junction, with the segmented density and model for the heavy chain only is shown in (**F).** The levers from the FH and BH approach the head-tail junction via a kink in the α-helix called the hook (bounded by two W residues, shown as sticks). The dashed blue circle indicates a region of unfolding of the ɑ-helix in the BH. (Movie S1B) **(G)** shows the head-tail junction, with both heavy and light chains included. Interacting residues between the RLC (FH) with the start of the coiled coil (Seg-1) are shown (K835:E32). No equivalent interactions are present for the RLC of the BH. (**H**) shows the density and model for the N-terminal region of the FH RLC from S19 to T10. Ionic interactions between R13 (FH RLC) and D55 and E51 (BH RLC) are shown. The phosphorylatable S19 and T18 residues are shown as spacefill. (**I)** shows an ionic interaction between the RLCs (K34: FH RLC with E35: BH RLC) just above the head tail junction.

### Head-tail junction: Heavy chains

We first focus on the head-tail junction (Fig. 1E, F), where the heavy chains (HCs) from each lever emerge from the motor domains, bending at W825 to form the hook (WQWW), before ending at the invariant proline (P836). The two HCs then associate to form the start of the coiled coil (Segment-1, Seg-1) at M847 (Fig 1E, F; Movie S1).

The structure of the free head (FH) and blocked head (BH) HCs following P386 are different. In the FH, there is very little unwinding of the ɑ-helix (2 residues to Q839). This may be accounted for by the interaction between E32 in the FH and K835 in the FH heavy chain (HC) (Fig. 1G). In contrast, in the BH HC, unwinding is more extensive (4 residues to E845) (dotted circle, Fig. 1F; Movie S1B) following P836, and the BH RLC does not interact with the BH HC (Fig. 1G). The unwinding of the BH HC additionally accommodates the bent path of the BH lever, allowing the BH HC to enter the start of the coiled coil (Seg-1) in register with that of the FH. The two HCs are in register at the start of the coiled coil (M847; Fig. 1F) and related by a 180° rotation around Seg-1 axis. Notably, the start of the coiled coil (Seg-1) takes a straight path down towards the FH as it emerges from the head-tail junction.

The overall structure of the head-tail junction in NM2A, while somewhat similar to that for full length shutdown SMM (Fig. S3 A-C), is markedly different to that reported for β-cardiac myosin (β-CM) heavy meromyosin (HMM) (Fig. S3D) (*18*). The BH levers take similar paths up to the hook for NM2A and β-CM, however the structure of the hook is twisted backwards and forms a more acute angle in β-CM compared to NM2A (Fig. S3E-F). In contrast, the FH levers take markedly different paths up to the hook (Fig. S3E, G). In β-CM, the FH lever is pulled closer towards the BH just after the converter, possibly as a result of the interaction between K611 in the BH motor domain and D143 in the FH ELC (*18*), absent in NM2A. Together with the altered hook structure of the β-CM FH and BH levers, the head-tail junction of β-CM is offset compared to NM2A, such that the Seg-1 of the coiled coil takes a path across the BH for β-CM (Fig. S3E, G) and not straight down between the two heads as for NM2A.

The lower resolution of the structure at the head-tail junction compared to other regions of the molecule (Fig. S1 E, G), is likely due to the variable nature of the head-tail junction (Fig. S3 H-J, Movie S2). This variability is likely important in allowing distortion in the heads when both are bound to actin in the same thin filament (*19*) and in optimizing mechanical performance (*20*). In shutdown myosin, it is likely important for the head-tail junction to accommodate distortions resulting from the formation of the interacting heads motif by the motor domains, the paths of the two levers up to the head-tail junction, and the emergence of the coiled-coil. Its variability mainly arises from the unfolding of the BH HC to allow for the twisting and translation of Seg-1 as it moves laterally to interact with Seg-3 and anteriorly/posteriorly to interact with the FH (Fig. S3H-J).

### Head-tail junction: Regulatory Light Chains

The N-terminal phosphorylation domain of the FH RLC (from S19 towards the N-terminus) encapsulates the interface between the RLCs (Fig. 1G, H, Movie S1C); a region we term the ‘mortar’ (*10*). Adjacent to the head-tail junction, we traced the path of the backbone of the FH RLC mortar to T10 improving on previous structures for smooth muscle myosin (SMM) (Fig. S4 A-D). This revealed an ionic interaction between R13 (FH RLC) with E51 and D55 (BH RLC), explaining why the N-terminal region of the FH RLC takes a path upwards between the two RLCs (Fig. 1H). Beyond T10, weaker unfilled density was consistent with the distal region adopting a range of positions. The two RLCs additionally interact outside of the phosphorylation domain (Fig. 1I), similar to that found for SMM (Fig S4 A-D).

The phosphorylatable serine, S19, is exposed on the surface of the mortar (Fig. 1H). Its phosphorylation would push S19 away from and weaken the RLC: RLC interface (Fig. S4E, F). T18, which can also be phosphorylated in cells resulting in diphosphorylated myosin (*21, 22*) is similarly exposed and available for phosphorylation (Fig. 1H). Using AlphaFold (*23, 24*), we built a structure of the kinase domain of myosin light chain kinase (MLCK) in complex with the RLC (Fig S3G). Placing this into the mortar in the shutdown structure, showed that S19 is accessible to and can interact with the kinase domain of MLCK with minimal steric clashes (Fig. S4H, I: Movie S1D). Phosphorylation of S19 in the mortar would increase the mobility and variability of the head-tail junction, which is likely to destabilize the shutdown state.

### Interactions of Segs-1 and -3 with the heads region and each other

After Seg-1 emerges straight down from the head-tail junction it continues down to interact with the FH loop 2 and HCM (hypertrophic cardiomyopathy) loop (Fig. 2A, B). In this region of the molecule, Seg-3 emerges from bend 2 and passes down behind the BH (Fig. 2A). Its initial path is likely guided by a set of interactions with the BH RLC and lever (Fig. 2C, D). Substitution of 2 of the charged residues involved (R814 and D1553 in NM2A and SMM) to non-polar (W) or polar (G,) residues respectively in β-CM) may contribute to the incomplete folding of β-CM (Fig. S6).

**Fig. 2.**
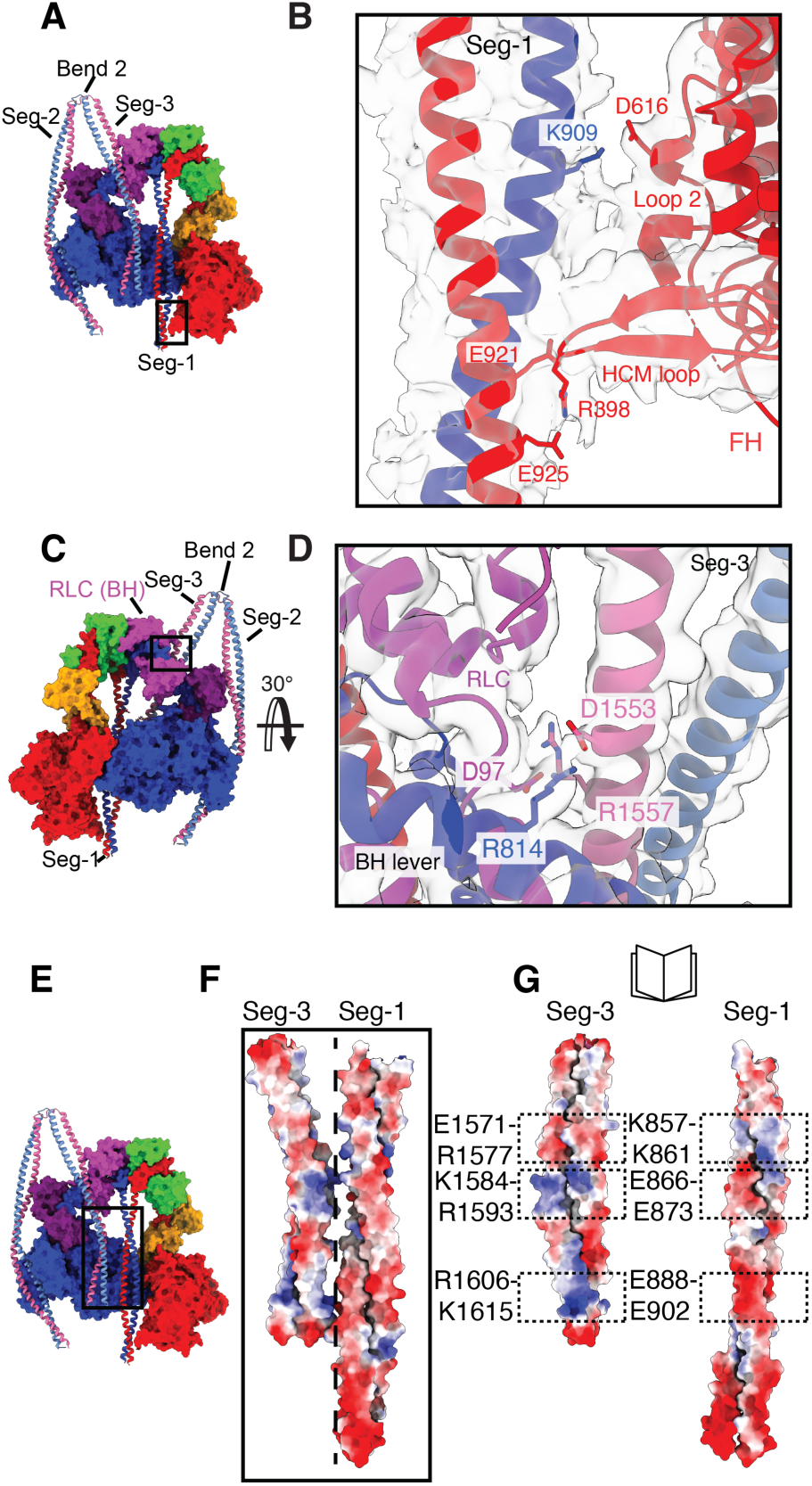
Interactions of Segs-1 and 2 with the motor domain and Segs-1 and -3 with each other. (**A)** Colored surface structure (face view) with the coiled coil shown as ribbons. The boxed region is expanded in (**B**) to show the ionic interactions between K909 (Seg-1) and D616 (loop 2) and E921 and E925 (Seg-1) with R398 (HCM loop). Zoned map at contour level 0.02. (**C**) Colored surface structure (back view) with coiled coil shown as ribbons. The boxed region is expanded in (**D**) to show interactions between Seg-3 and both the RLC and lever of the BH. (D1553:R814, D97 (BH RLC): R1557). Zoned map at contour level 0.02 (**E**) Colored surface structure (front view), with coiled-coils shown as ribbons to show the positions of Seg-1 and Seg-3 as they pass across the blocked head (boxed region). Expanding the boxed region in (**E**) in (**F**) shows the multiple potential ionic interactions between Seg-3 (BH) and Seg-1 (FH), each colored by electrostatic potential. (**G**) The interface between Seg-3 and Seg-1 is opened up (as a book) to display the electrostatic seam between the two pairs of CCs. Dashed boxes show patches of opposite charge between Seg-1 and -3.

Close to the motor domains, Segs-1 and -3 begin to strongly interact through a series of multiple ionic interactions (Fig. 2E-G) such that there is a tight association between these two regions of coiled coil. Seg-1 does not run across the BH as it does in β-CM (Fig. S5A-C). While Seg-3 appears to do so, it only makes 2 interactions with the BH (Fig. S5 A, C). The equivalent interaction for the first of these interactions (R442 (loop O): E1593 (Seg-3)) is also found in SMM (Fig. S5D) and in β-CM (K450 (loop O): E875 (Seg-1). The second interaction (K656 (helix W): D1600 (Seg-3)), not found in SMM, is equivalent to the T660 (helix W): Q892 (Seg-1) interaction in β-CM. In β-CM, Seg-1 in the heavy meromyosin (HMM) shutdown structure goes on to make multiple interactions with the BH (Fig. S5B, E), as part of the so-called mesa trail (*25*). In this case residues in and downstream of Ring-1 in Seg-1 interact with residues in the motor domain. Similar interactions are also found in shutdown molecules in the cardiac myosin filament (*26*). Many of the residues involved in these interactions are conserved in NM2A and SMM (Fig. S6), suggesting Seg-1 could interact with the blocked head in a similar way to β-CM in these myosins. However, in the full length molecule, Seg-3 prevents Seg-1 from accessing the BH in NM2A and SMM (*27*). The small number of contacts between Seg-3 and the BH (Fig. S5C, D), together with the tight association between Segs-1 and -3 in NM2A, suggest that the interactions between the coiled coils dictate the path of Seg-1 and -3 in this region of the molecule.

Overall, the straight path of Seg-1 and its interactions with loop 2 of both blocked and free head, together with the strong interactions between Seg-1 and -3, and the interactions of Seg-3 with the BH RLC and lever help to explain the path of Seg-1 and-3 in the heads region.

### Seg-2 and its interactions with the BH, Bend 2 and the Latch

Below the myosin heads, Seg-1 continues down to bend 1. Bend 1 was identified as a region of unwinding around residue A1156, using a combination of crosslinking mass spectrometry data and AlphaFold modelling of the missing coiled segments (the density in this region is too weak to fit the structure (Fig. S7 and see methods)).

The coiled coil then returns back towards the BH as Seg-2. It first makes key interactions with residues from the N-terminal SH3-like fold and the SH1-helix of the BH motor domain, sitting within a small groove formed by these two regions (Fig. 3A-C), similar to our findings for SMM (*10*). The improved resolution of our NM2A structure allows us to confidently identify key interacting residues between the motor and Seg-2 in this region. Missense mutations in any one of 8 out of the 11 interacting residues, have been linked to MYH9 disease highlighting the importance of this region of the structure in stabilizing the shutdown state (*4, 28*). In agreement, we previously reported that missense mutations in NM2A in one of these key residues (D1424), commonly mutated in MYH9 disease (*28*), destabilizes the shutdown state (*4*). In our NM2A structure, we additionally resolved the complete N-terminal region upstream of the SH3-like fold of both blocked and free heads (Fig. S8), revealing how this region of the structure lies across the motor domain. MYH9-disease related mutations in this region likely affect the subsequent positioning of the SH3-like fold, destabilizing its interaction with Seg-2.

**Fig. 3:**
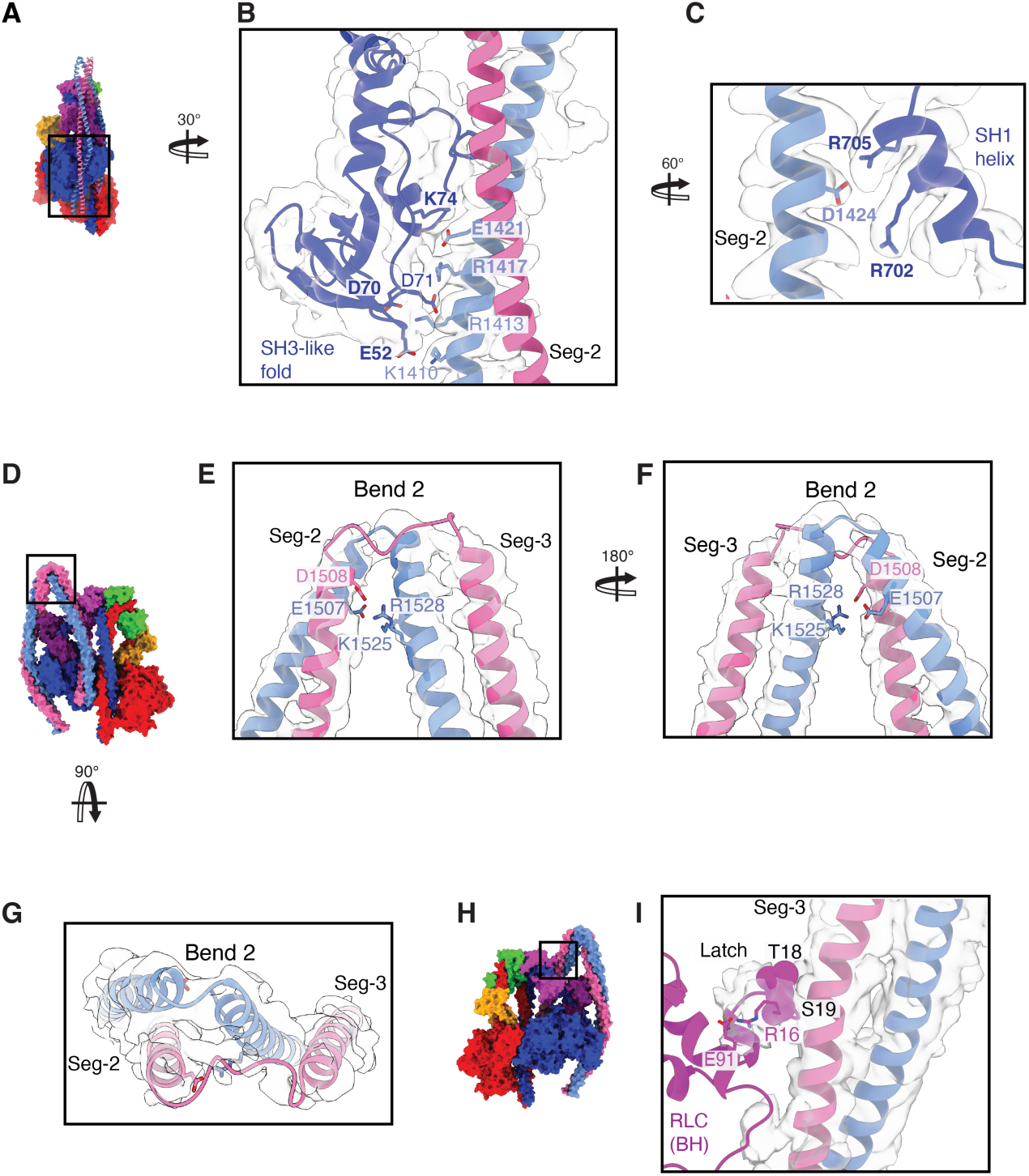
Interactions between Seg-2 and the BH motor domain, formation of Bend 2 and the N-terminal region of the BH RLC (Latch). (**A**) Side view of the motor domain of the blocked head (colored surface structure), showing how Seg-2 of the coiled coil sits in a groove within the BH motor. Two views of the boxed region in **A** show the ionic interactions between residues in the N-terminal SH3-like fold (E52, D70, D71 and K74) with Seg-2 (K1410, R1413, R1417 and E1421) (**B,** zoned map at contour level 0.02), and the ionic interactions of specific residues (R701 and R704) in the SH1 helix of the motor domain (BH) with Seg-2 (D1424) (**C,** zoned map at contour level 0.1). Residues shown in bold type are mutated in MYH9-related disease (**D**) Face view of bend 2 (colored surface structure). the boxed region is shown expanded in face view (**E**), back view (**F**) and top view (**G**) to show the local unwinding of the coiled coil (zoned map at contour level 0.1.) and the ionic interactions that stabilize this bend. (**H**) Back view (colored surface structure). (**I)** shows the boxed region in **H**, to show residues S19 to R16 in the N-terminal region of the BH RLC (latch) and their proximity to Seg-3. The phosphorylatable S19 and T18 residues are shown as spheres. Zoned map at contour level 0.02.

The interactions we observed between Seg-2 and the BH are similar to those that we previously suggested would be present in SMM from our pseudoatomic model at lower resolution (*10*) (Fig. S9A-C, E-G). However, a more recent SMM structure (7MF3) does not appear to show similar interactions (Fig. S9 D, H). To try to understand this, we compared our coiled-coil structure in NM2A to that in SMM (7MF3: Fig. S10) and noticed that the coiled coil in 7MF3 goes out of register in Seg-2 (Fig. S10A, B)(*9*). A transverse view through the coiled coil in a region of Seg-2 from NM2A (residues 1408-1411) shows how the two chains are in register, with sufficient density to fit the side chains for the *a* (Y1408) and *d* (L1411) residues, demonstrating that they are embedded within the hydrophobic seam (Fig S10C). This is not the case in the equivalent region of 7MF3; both *a* (Y1421) and *d* (L1424) residues are not in register, and neither are embedded within the hydrophobic seam (Fig. S10D). Thus, it is likely that the coiled coil in 7MF3 was not built correctly and cannot therefore be used to identify interactions between Seg-2 and Seg-3 and the rest of the molecule.

As Seg-2 continues towards bend 2, it first makes 2 ionic interactions with the ELC (Fig. S11), similar to our previous findings for SMM. These interactions may bias the path of Seg-2 towards bend 2. The two residues in Seg-2 (E1472 and R1482) that interact with the ELC are not conserved in Seg-2 of β-CM (Fig. S11C, D; Fig S11). This could explain why, in isolated molecules of β-CM, the coiled coil may weakly interact with the motor domain in the region of the groove (*29*) but then does not make any further interactions with the motor and bend 2 fails to form.

The resolution of our NM2A structure reveals that bend 2 in NM2A, is facilitated by a small unwinding of the coiled coil, rather than a bend at a single residue (Fig.3 D-G). The unwinding we observe in NM2A is similar to that for SMM, although we attributed bend 2 to a single glycine residue (*10*). In NM2A, 10 residues in the FH ɑ-helix are unwound at bend 2 (M1510 to S1519) and G1516 (equivalent to the single glycine residue identified as bend 2 in SMM) is within this region. However, only 2 residues in the BH ɑ-helix (K1513 to D1514) are unwound. Ionic interactions between Seg-2 and Seg-3 help to stabilize bend-2 (Fig.3 E, F).

As Seg-3 passes back down to interact with Seg-1 after bend 2, it is likely to interact with the N-terminal region of the FH RLC, that we previously termed the latch. We resolved the path of the backbone of the latch up to R16, an improvement on our previous SMM structure (Fig. S12 A-D) (*9–11*). The unfilled density beyond this was weaker, suggesting that this distal region can adopt various conformations, all close to Seg-3. Phosphorylation of S19 in the latch would cause side-chain clashes and repulsion by negatively charged residues in Seg-3 (Fig. S12 E-G) releasing the latch from its shutdown position. Docking the kinase domain from MLCK showed that its access to S19 is sterically blocked by Seg-3 (Fig. S12 H, I) in contrast to our finding for the mortar. This suggests that S19 in the mortar (FH RLC) is phosphorylated first, and subsequent disruption of the shutdown state then allows S19 in the latch to be phosphorylated, as we previously speculated for SMM (*10*). This contrasts with a previous report that suggests that S19 in the mortar is less accessible, but which did not explore the ability of MLCK to bind to the latch and mortar (*9*).

### Full Length shut-down NM2A

We additionally solved the structure of an extended region of shutdown NM2A to a global resolution of 6.3 Å (Fig. 4A-D, Fig. S1). This extends the structure to about 2/3^rd^ of the way along the coiled-coil tails. The high mobility of the distal region (Movie S3) prevented us from solving the entire structure from the density map. However, combining cross-linking mass spectrometry data with AlphaFold modelling (see methods) enabled us to build in the missing coiled coil segments outside the density map. This structure of the full-length molecule shows how the three segments of the coiled coil come together and interact in NM2A (Fig. 4E, F), as we suggested for SMM, to form an untwisted ribbon structure (*10*). Seg-1 lies between Seg-2 and Seg-3. A coulombic plot shows the alternating positive and negatively charged regions that enable the interaction between Segs-1 to 3 in this region (Fig. 4E, F).

**Fig. 4.**
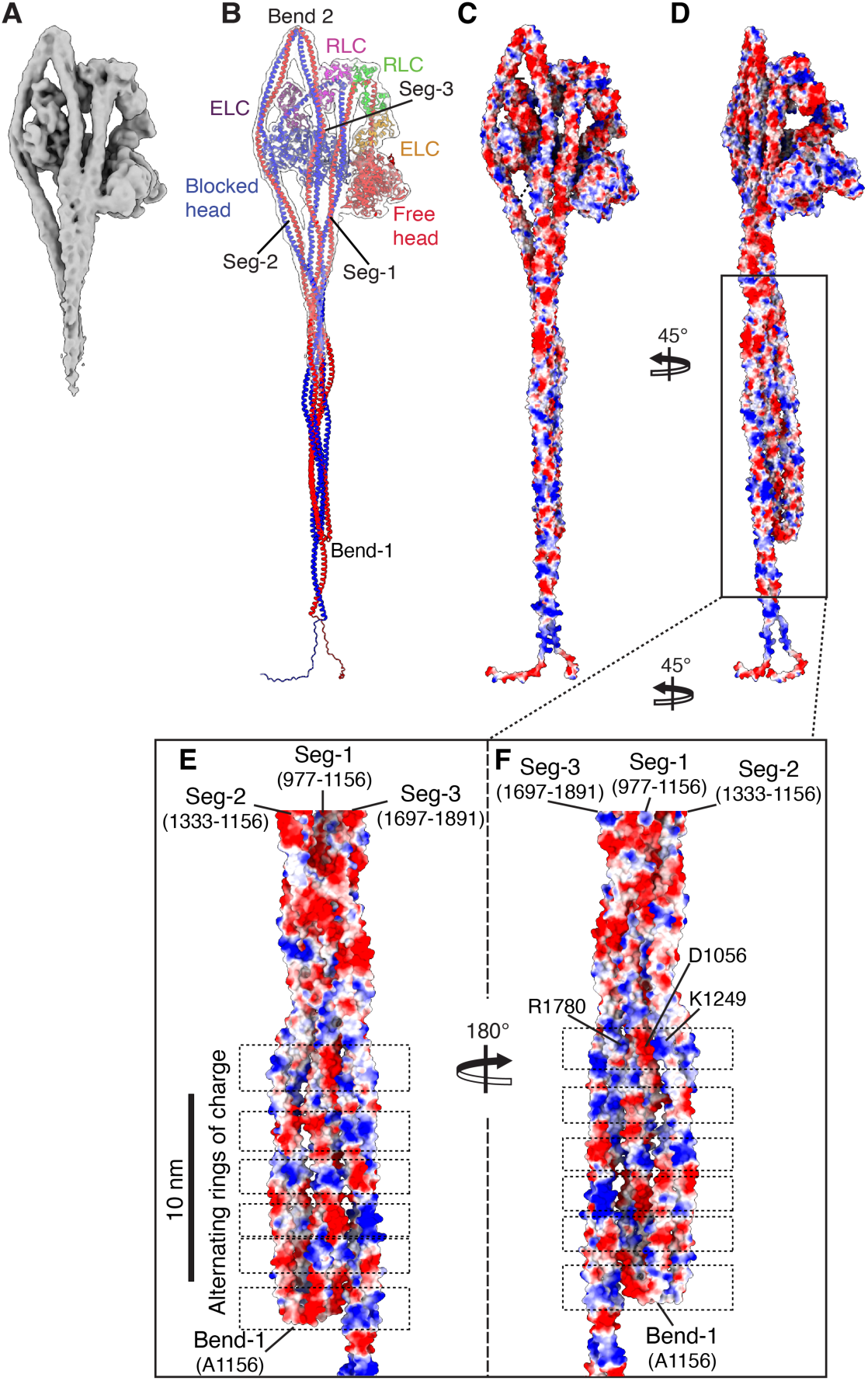
Structure of the whole 10S complex of NM2A and the flat ribbon arrangement of its coiled-coil tails. (**A).** Cryo-electron density map of the full-length molecule (front view). (**B**) the structure of whole molecule fit into the density map. The structure has been colored by chain in the same way throughout: Heavy chain for the motor, lever and the first part of the coiled coil (segment 1) shown in red for the free head, and blue for the blocked head. (chain assignment as (*10*) for SMM) ELC: dark purple for blocked head and orange for free head. RLC: green for free head and magenta for blocked head. **(C)** Surface representation (face view) and (D) side view of the structure in **B**, colored by electrostatic potential. **(E)** and **(F)** show expanded views of the boxed region in **D** showing alternating rings of charge which keep the segments sandwiched together.

### Model for activation of NM2A by RLC phosphorylation

If S19 in the mortar is phosphorylated first, as our data seems to indicate, we can build a picture of how the shutdown molecule would become destabilized when the RLCs are phosphorylated (Fig. 5). Phosphorylation of FH RLC S19 (Fig. 5A) would destabilize the mortar, which in turn would destabilize RLC-RLC interactions at the head-tail junction (Fig. 5B). This would lead to an increased mobility of the head-tail junction, and thus Seg-1 and 3, which are closely associated (Fig. 5C). This in turn would increase mobility of Seg-2, pull Seg-2 out of the groove N-terminal formed by the SH3-like fold and the SH1-helix and away from the ELC leading to the release of Seg-2 from the IHM. The increased mobility of all segments of the tail would expose the latch, allowing access of MLCK to S19, and its phosphorylation (Fig. 5D). With the latch phosphorylated, Seg-3 would be released from the tail ribbon (Fig. 5E), leading to separation of all three tail segments and dissolution of the IHM resulting in an active open molecule (Fig. 5F).

**Fig 5.**
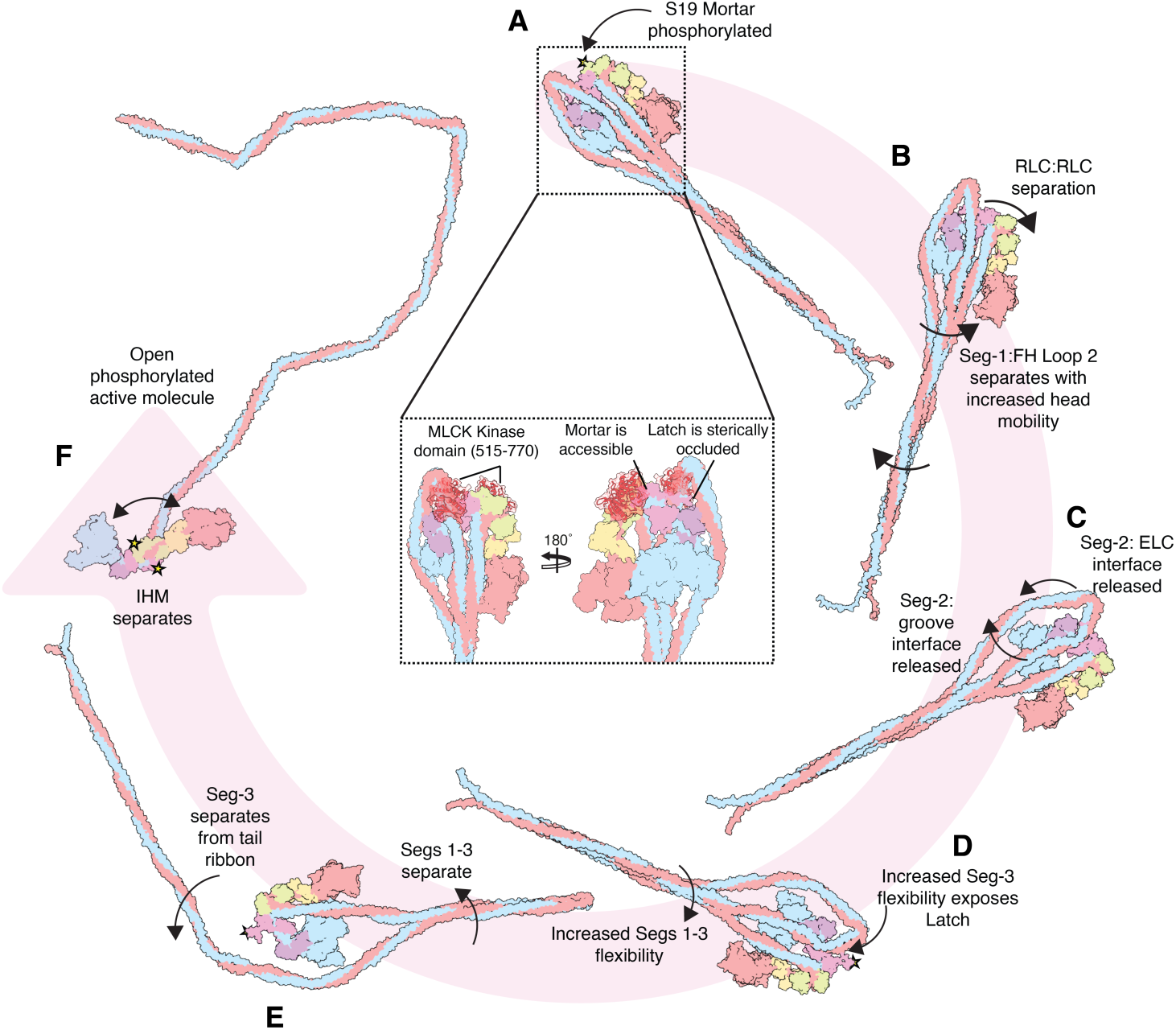
Possible sequence of 10S to 6S transition. (**A**) Intact 10S molecule is phosphorylated at the mortar (phosphorylation site indicated by the gold star) Insets show the different accessibility of the MLCK Kinase domain to the RLC N-terminal extensions of FH and BH (**B**) Destabilization of the RLC-RLC interface increases the mobility of Seg-1 at the head-tail junction, allowing Seg-1 to separate from the FH. (**C**) Increased mobility of the tail segments allows Seg-2 to be released from the SH3 groove and ELC interface (**D**) the increased mobility of all 3 segments of the tail exposes the latch to become accessible to MLCK and then phosphorylated (gold star indicates phosphorylation site) (**E**) With the latch phosphorylated, Seg-3 is released from Seg-1 and then the **(F)** molecule is completely unfolded and active.

In conclusion, the shutdown NM2A structures presented here reveal how NM2A is shutdown in its dephosphorylated state, the key residues that stabilize this state, and how phosphorylation activates the molecule. The roles of key residues in stabilizing this shutdown state bring new understanding into how mutations could destabilize it, leading to disease.

## Author contributions

MP designed the project, obtained the funding, supervised and led project administration. D-CM and GC provided resources, carried out the investigation, curated and analyzed the data and validated the results. MP, D-CM and GC visualised the data and wrote, reviewed and edited the paper. We thank staff at the Astbury Biostructure Laboratory for their support and members of the CryoEM community at Leeds for their help and guidance. We specifically thank Peter Knight and Charlie Scarff for comments on previous drafts of the manuscript.

## Data and materials availability

the structure of the heads region of shutdown NM2A has been deposited in the PDB: (PDB ID 9SYU: EMD-55354), and that for the full-length structure (PDB ID 9SZR: EMD-55382).

## Supplementary Materials for

### Materials and Methods

#### Protein expression and purification

NM2A protein was obtained by co-expression of FLAG-tagged human NM2A heavy chain (MYH9 coding sequence: NCBI: NG_011884.2) with bovine non-muscle myosin RLC (MYL9: 95% identical in amino acid sequence to human) and chicken non-muscle myosin ELC (MYL6: 90% identical in amino acid sequence to human) as reported previously (32), using Sf9 cells. The protein was purified using M2 FLAG affinity gel (Sigma) as described previously (4, 30). NM2A expressed in this way is not phosphorylated (31).

To form the shutdown state, Mg.ATP was added to the purified NM2A, before dilution into a low ionic strength solution. Final conditions were 140 mM KCl, 10 mM MOPS pH 7.2, 0.1 mM EGTA, 2 mM MgCl_2_,1 mM ATP. 1 mM Bissulfosuccinimidyl suberate (ThermoFisher) was used to cross-link 1 μM 10S particles for 30 min at 25 °C and then quenched with the addition of pH 8.0 Tris buffer to a final concentration of 100 mM.

#### Grid preparation and cryo-EM data acquisition

Grids were prepared with a Vitrobot Mark IV (Thermo Fisher), applying 3 μl of NM2A (0.6 μM) to UltrAuFoil R1.2/1.3 Gold 300 mesh grids (Agar Scientific, UK) glow discharged for 90 s before use (PELCO easiGlow discharge unit, Ted Pella Inc). Grids were blotted with Whatman no. 42 Ashless filter paper (Agar Scientific, UK) for 3 s at force 6, at a temperature of 8 °C and 100% humidity, and flash-frozen in liquid ethane. Data were recorded on a FEI Titan Krios (Astbury Biostructure Laboratory, University of Leeds) operating at 300 kV equipped with a TFS Falcon 4i direct electron detector and a Bio-quantum energy filter (Gatan, CA, USA). Micrographs were recorded with the EPU automated acquisition software at 96,000× nominal magnification, giving a final object sampling of 0.82 Å/pixel, a total dose of 37 e^−^/Å^2^, and a target defocus range between −0.9 and −3.0 μm (4). We acquired 27,549 micrographs in two microscope sessions using the same collection parameters (Table S1).

#### Cryo-EM image processing

Image processing was carried out using cryoSPARC v4.6.0 (33). Drift-corrected averages of each movie were created using the Patch Based Motion Correction in cryoSPARC and the defocus of each was determined using Patch CTF (34). Micrographs were curated to exclude micrographs with excessive ice thickness, poor CTF fit, and large full-frame motion. Particle picking was performed using a combination of Blob Picker functionality and the Topaz wrapper in cryoSPARC (35, 36). Particles were extracted in a box size of 480 × 480 pixels, centered on the interacting heads motif (IHM) region and were classified using 2D classification in cryoSPARC. Classes that contained features reflective of shutdown myosin structure were selected and taken forward for further classification in cryoSPARC. Four initial 3D volumes were produced using ab-initio reconstruction in cryoSPARC. This model was refined with several rounds of heterogeneous and non-uniform refinement. The final particle stack contained 349,374 particles from 20,755 micrographs. Reference-Based Motion Correction and a final 3D refinement yielded a map for the heads region of the full-length molecules that had a 3.0 Å global resolution determined using the gold standard Fourier shell correlation (FSC) reported to FSC = 0.143 (Fig. S1). The EM density map was sharpened using a negative *B*-factor that was automatically determined in cryoSPARC using a Guinier plot. Local resolution was estimated using cryoSPARC.

To obtain a structure of the full length NM2A molecule, particles used to generate the 3.0 Å global resolution map of the NM2A heads region in cryoSPARC were re-extracted in a box size of 1024 × 1024 (binned twofold into a 512 × 512 box). An initial 2D classification was performed. Good particles were first classified in 2D, followed by ab-initio reconstruction and non-uniform refinement in cryoSPARC. The final global resolution of the NM2A shutdown whole-molecule map was 6.3 Å (from 15,167 particles), determined using the gold standard Fourier shell correlation (FSC) reported to FSC = 0.143 using cryoSPARC. Figures and videos were generated in ChimeraX (37) or Chimera (38).

#### Model building and refinement

To interpret the 3.0 Å map for the IHM region, we created an atomic model for the myosin heads, using ModelAngelo (39) to automatically build into the map, using the sequences of the heavy chain, regulatory light chain (RLC), and essential light chain (ELC). Mg^2+^ was present in both the nucleotide binding pocket and in the RLC. Unbuilt portions were performed manually using the SMM structure (10) as a homology model and then using flexible fitting using ISOLDE (40). Coot (41) was used to address bond length and geometry issues introduced by molecular dynamics simulation of model building using ISOLDE.

To generate a full-length NM2A model, a full-length SMM homology model based on the whole SMM structure was used in ISOLDE to build into the extended map. Chains C, D, E, and F were built to the same extent as in the heads-only model. Segs-1, 2, and 3 were built into density from residues 847-1072, 1235-1516, and 1517-1794, respectively. AlphaFold3 was used to generate three model coiled-coil segments for the missing tail sequences, which each contained 14 residues that overlapped the tail sequences of Segs-1, 2 and 3 of our pseudo-atomic model of the IHM region. These models were superposed onto the existing coiled-coil tail segments and then the ‘fit selection only’ flag was used in rigid body fitting as described for our 9 Å map of the whole SmM shutdown molecule (10). This allowed the introduction of the bends into the coiled-coil tail and for them to lie in plane as seen in the density map. With Segs-1 and -2 lying antiparallel to one another, the point at which their sequences coincided was found to occur at A1169, so bend 1 was created at this point. The chains were joined and duplicated sequence was deleted followed by manual refinement in Coot. The junction made between Segs-1 and 2 at bend 1 implies that in Segs-2 and 3, chain G is a continuation of chain A and chain H is a continuation of chain B. Cross-linking distance restraints were used together with ISOLDE to validate the final configuration. As the entirety of each heavy chain can be built, the full-length model follows the same chain nomenclature as above but with chains G and H being built as a continuation of chains A and B, respectively.

The structure of the bovine RLC (MYL9, residues 1–173) in complex with the human myosin light chain kinase 3 (MYLK3, kinase domain 491–746), which we term the RLC_MLCK complex, was generated using the AlphaFold3 server (https://alphafoldserver.com/) (43). The top scoring model was aligned using residues 17-23 of the RLC N-terminal extension in the RLC_MLCK3 complex onto each RLC chain (chains E and F; residues 17 – 23) of our 10S structure using the ‘Matchmaker’ command within USCF ChimeraX. With the RLC_MLCK complex docked into place, the RLC chain from the RLC_MLCK complex was deleted to yield the MLCK kinase domain in complex with both the free-head and blocked-head RLC chains of our 10S structure.

#### Crosslinking Mass Spectrometry

Cross-linking mass spectrometry was performed as previously described (44). Briefly, crosslinked samples were reduced (20 mM Dithiothreitol (DTT)), alkylated (40 mM Indole-3-acetic acid (IAA)), acidified, and digested on S-Trap columns (ProtiFi) with trypsin at 47 °C for 90 min. Peptides were desalted, concentrated, and reconstituted in 0.1% formic acid. Liquid Chromatography-Mass Spectrometry (LC-MS/MS) was carried out on an Orbitrap Eclipse Tribrid with a Vanquish Neo UHPLC (Ultra-high performance liquid chromatography), using a PepMap C18 trap and EASY-Spray C18 analytical column with a 95-min 2–50% acetonitrile gradient. Data-dependent acquisition was used (MS1: 120k resolution, 380–1450 m/z; MS2: HCD, 60k resolution, dynamic exclusion 30 s). Data were analyzed in Proteome Discoverer v3.0 (XlinkX) and Merox v2.0.1.4 with a custom FASTA (precursor 5 ppm, fragment 10 ppm, trypsin with 2 miscleavages, carbamidomethyl Cys fixed, Met oxidation and N/Q deamidation variable, FDR < 1%, score > 50).

**Fig. S1.**
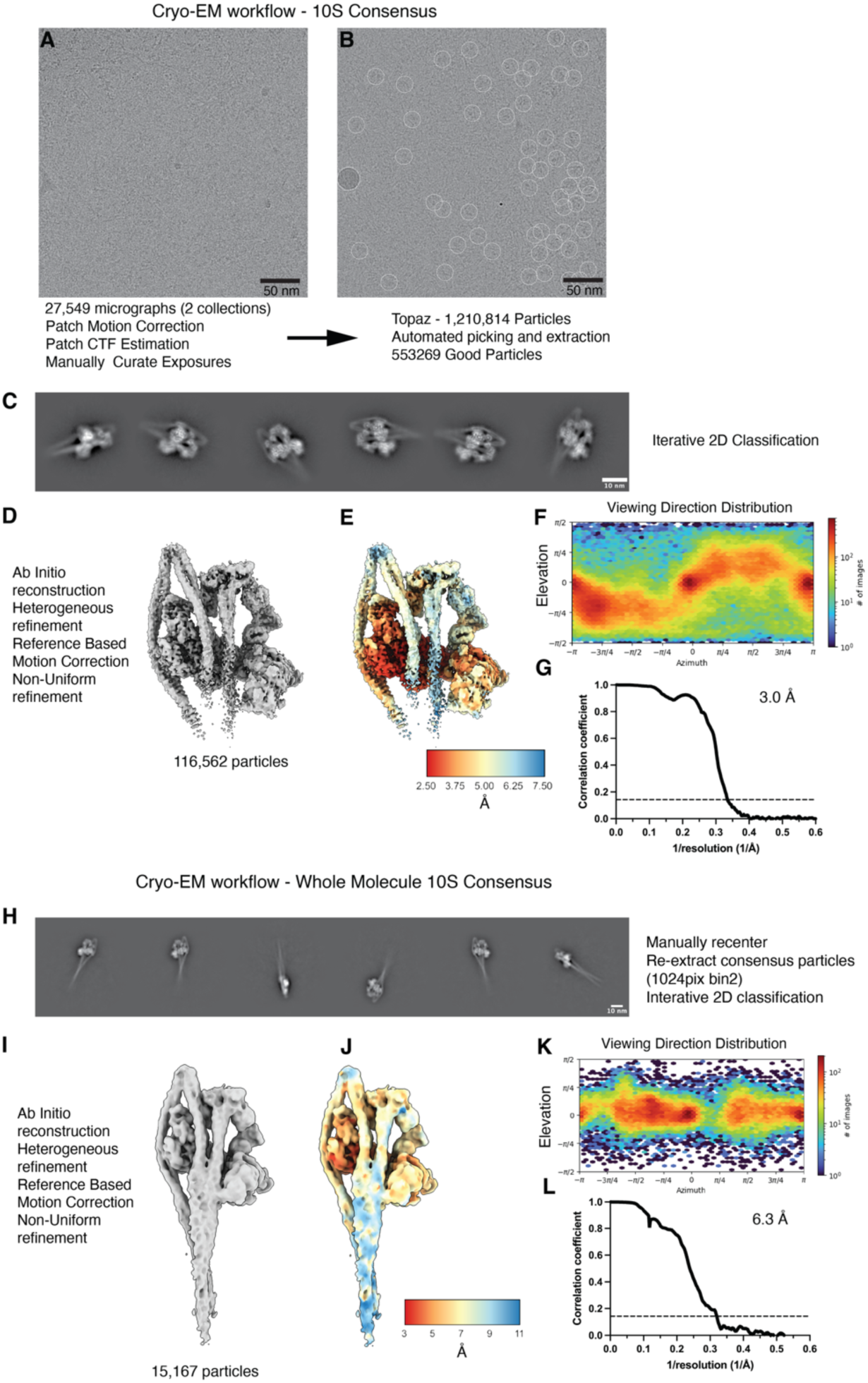
NM2A Processing Workflow. (**A**) Representative micrograph of shutdown NM2A molecules. (**B**) Representative micrograph with particle picks. (**C**) Representative 2D classes (Scale bar 10nm). (**D**) Final NM2A shutdown heads region cryo-EM map (deposited as EMD-55354). (**E**) Local resolution of map shown in **D**. (**F**) Angular distribution of particles in the heads region 3D reconstruction. (**G**) FSC curve of the heads region reconstruction, illustrating 3.0 Å resolution at 0.143 FSC. (**H**) Representative 2D classes of the whole molecule. Particles were extracted in a 1024 pixel box and binned twofold. (Scale bar 10nm) (**I**) Final NM2A shutdown whole molecule cryo-EM map (deposited as EMD-55382). (**J**) Local resolution of map shown in **I**. (**K**) Angular distribution of particles contributing to the whole molecule 3D reconstruction. (**L**) FSC curve of the whole molecule reconstruction, illustrating 6.3 Å resolution at 0.143 FSC.

**Fig. S2.**
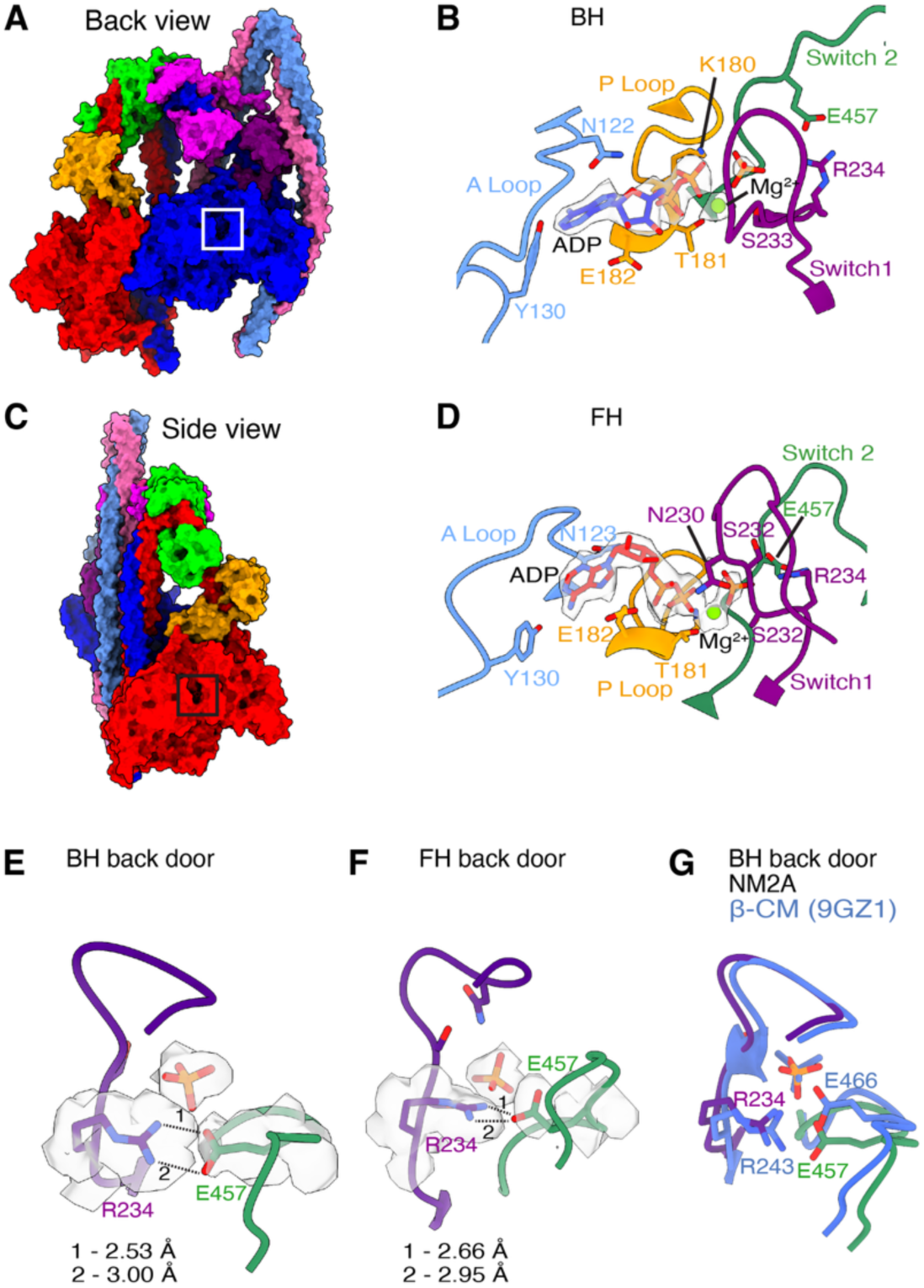
The nucleotide binding pocket in shutdown NM2A. (**A**) Colored surface view of the heads region (back view) to show the location of the blocked head (BH) nucleotide pocket. (**B**) Zoomed view of region boxed in **A**. Side chains of interacting residues in the A Loop, P Loop, Switch 1 and Switch 2 are indicated. ADP.Pi and Mg^2+^ are fit to segmented density. (**C**). Colored surface view of the heads region (side view) to show the location of the Free Head (FH) nucleotide pocket. (D) Zoomed view of region boxed in **C,** with equivalent interacting loops and residues indicated as in **B**. (**E-G**) Salt bridge blocking Pi release and corresponding bond lengths in the BH and FH back door. (**G**) Alignment of NM2A and β-CM (PDB:9GZ1) BH Back Door.

**Fig S3:**
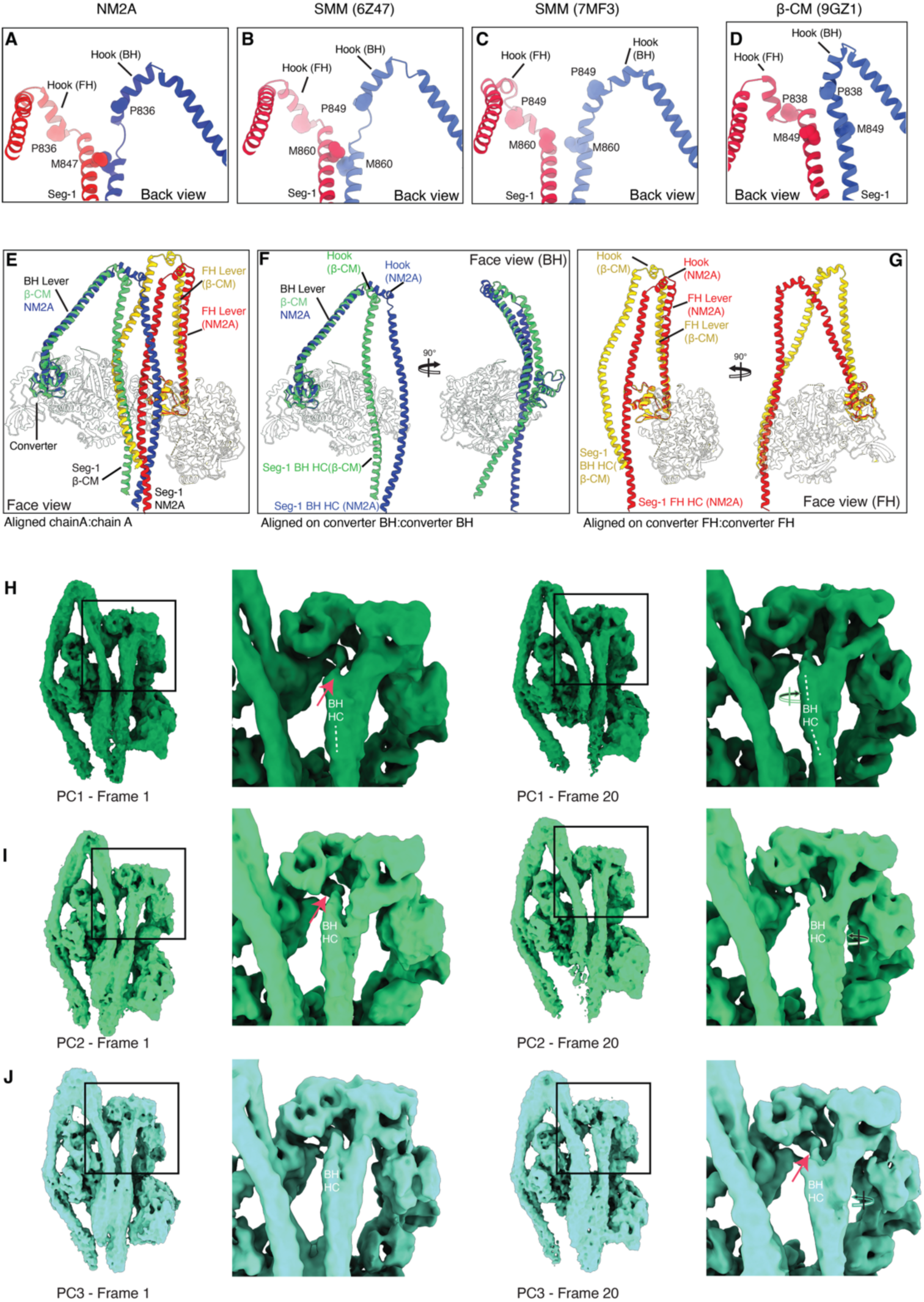
Comparison of β-CM and NM2A levers and Head-Tail junction. (**A**) Back view of the Head-Tail junction of NM2A. Light chains are hidden and the side chains of the invariant proline that marks the start of the Head-Tail junction conserved methionine are shown as spheres. (**B-D**) Same representation as in A for 6Z47 (*10*), 7MF3 (*9*), and 9GZ1 (*18*) respectively. (**E**) Alignment of shutdown β-CM HMM and shutdown NM2A whole molecule HCs in the region of the heads. Light chains are hidden and heavy chain (HC) residues before the converter are grayed out for clarity. (**F**) Alignment of β-CM and NM2A BH HCs. Alignment was performed on the converter to emphasize the divergent path of the hook and Seg-1. (**G**) Alignment of β-CM and NM2A FH HCs. Alignment was performed on the converter to emphasize the divergent path of the LCD, hook, and S1. **H, I and J** show the first and last frame of the 3 component Principal Component Analysis of the 3D Variability Analysis performed in CryoSparc. Arrows indicate the twisting and lateral movement of Seg-1 as it emerges from the head-tail junction across the population of conformations. (See Supplemental movie 2).

**Fig. S4.**
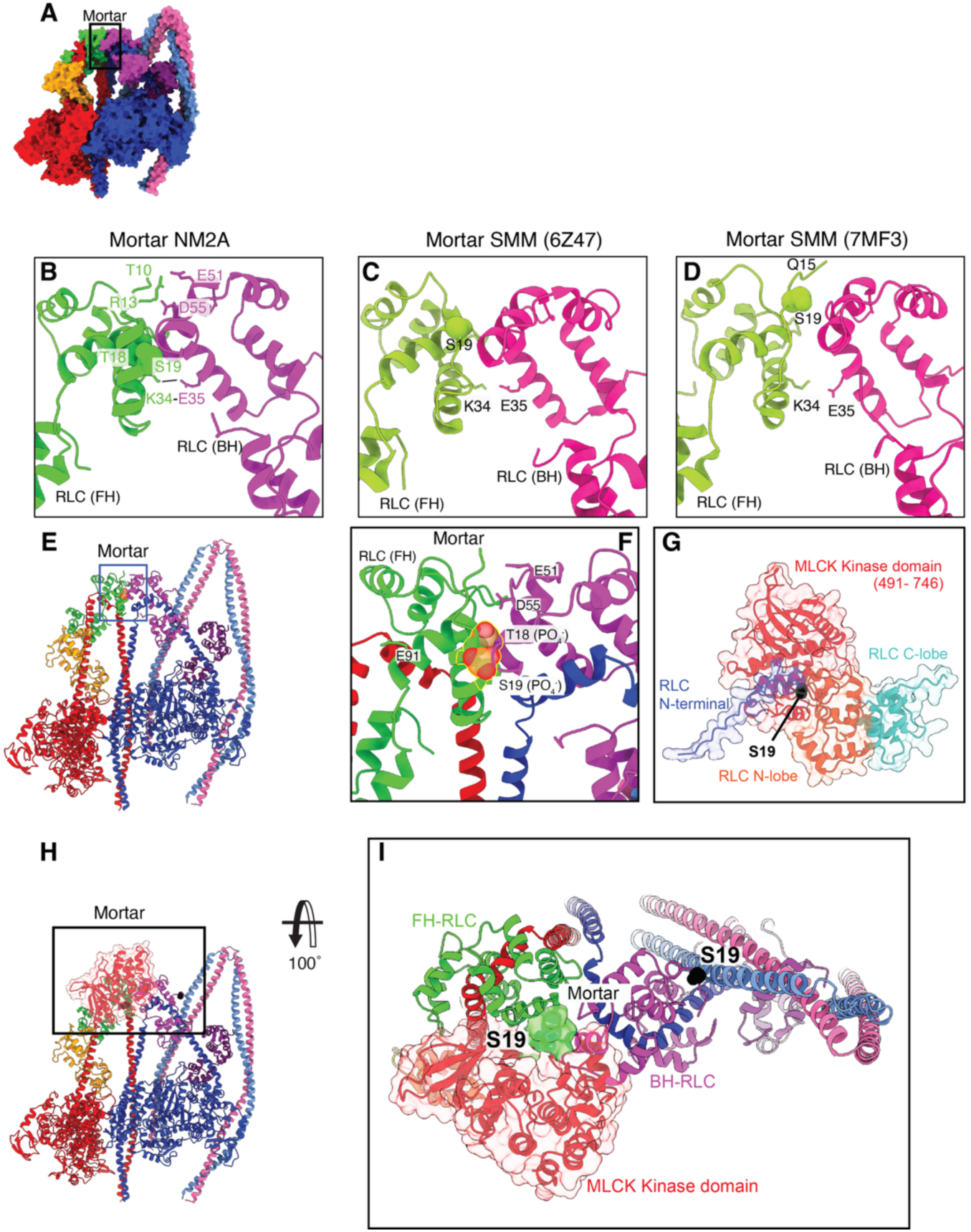
Comparison of Mortar for NM2A and SMM and its phosphorylation by MLCK. **(A)** Colored surface view of the heads region (back view) to show the location of the FH RLC N-terminal extension (Mortar) **(B)** Zoomed view of region boxed in **A** showing RLCs only (HCs omitted). Side chains of interacting residues between the two RLCs are shown. Side chains of the phosphorylatable S19 and T18 residues are shown as spheres. **(C, D).** Equivalent views (as in **B**) for SMM (6Z47 (*10*) and 7MF3 (*9*) (**E)** Ribbon structure of the modelled phosphorylated molecule **(F)** A zoomed in view of the phosphorylated mortar. Phospho-S19 and Phospho-T18 residues are shown as spheres, with side chains of clashing residues in the mortar interface shown as sticks **(G)** The top scoring AlphaFold model of the MLCK Kinase domain (491–746) in complex with Bovine RLC (RLC_MLCK), showing the N-terminal extension of the RLC embedded in a groove of the kinase domain (S19 shown in black). **(H)**To place the kinase domain into the shutdown structure, the N-terminal extension of the RLC kinase: RLC complex (residues 17 – 23) was aligned onto the RLC of the BH. **I** shows boxed region in **H** as a surface fill representation of the BH N-terminal extension (green) with the kinase domain (red). The kinase makes minimal contacts with the rest of the RLC.

**Fig. S5:**
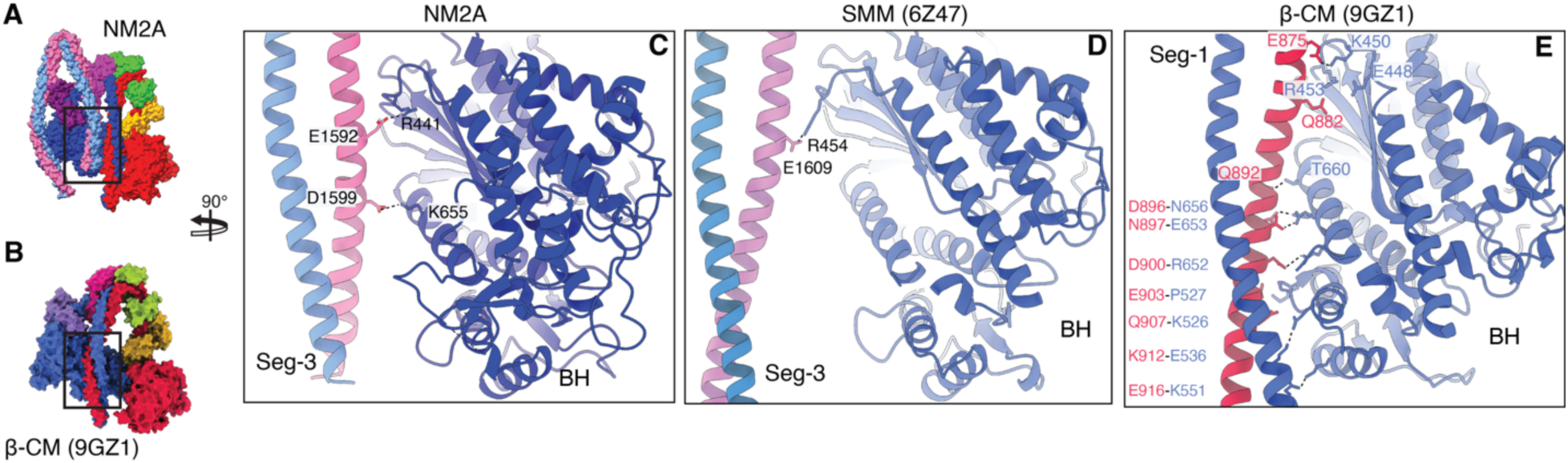
Paths of Seg-1 and Seg-3 in β-CM and NM2A/SMM, respectively. (**A**) Colored surface view of the heads region of NM2A (face view) showing Seg-3 and its position close to the BH (**B**) Colored surface view of the heads region of β-CM (face view: 9ZG1 (*18*)) to show the location of Seg-1 (**C**) Zoomed and rotated view of the region of NM2A boxed in **A**. Side chain interactions between Seg-3 and the blocked head are shown. (Numbering excludes methionine as residue 1). No other interacting residues were present. (**D**) Equivalent view for SMM (6Z47 (*9*) as in C. (**E)** Zoomed and rotated view of the region of β-CM boxed in **B.** Side chains of interacting residues between Seg-1 and the mesa trail are shown (interacting residues are indicated on the LHS, as CC – BH). The view shown is the equivalent representation as in **C, D** for the alignment of β-CM (9GZ1) with NM2A and 6Z47, with Seg-1 occupying a similar space to Seg-3. The mesa trail in β-CM is extensive (18) and only some of the interactions reported are shown. Residues D896, N897, D900 and E903 (β-CM) are all in a region of acidic charge termed Ring-1 (see alignment in Fig. S6).

**Fig. S6:**
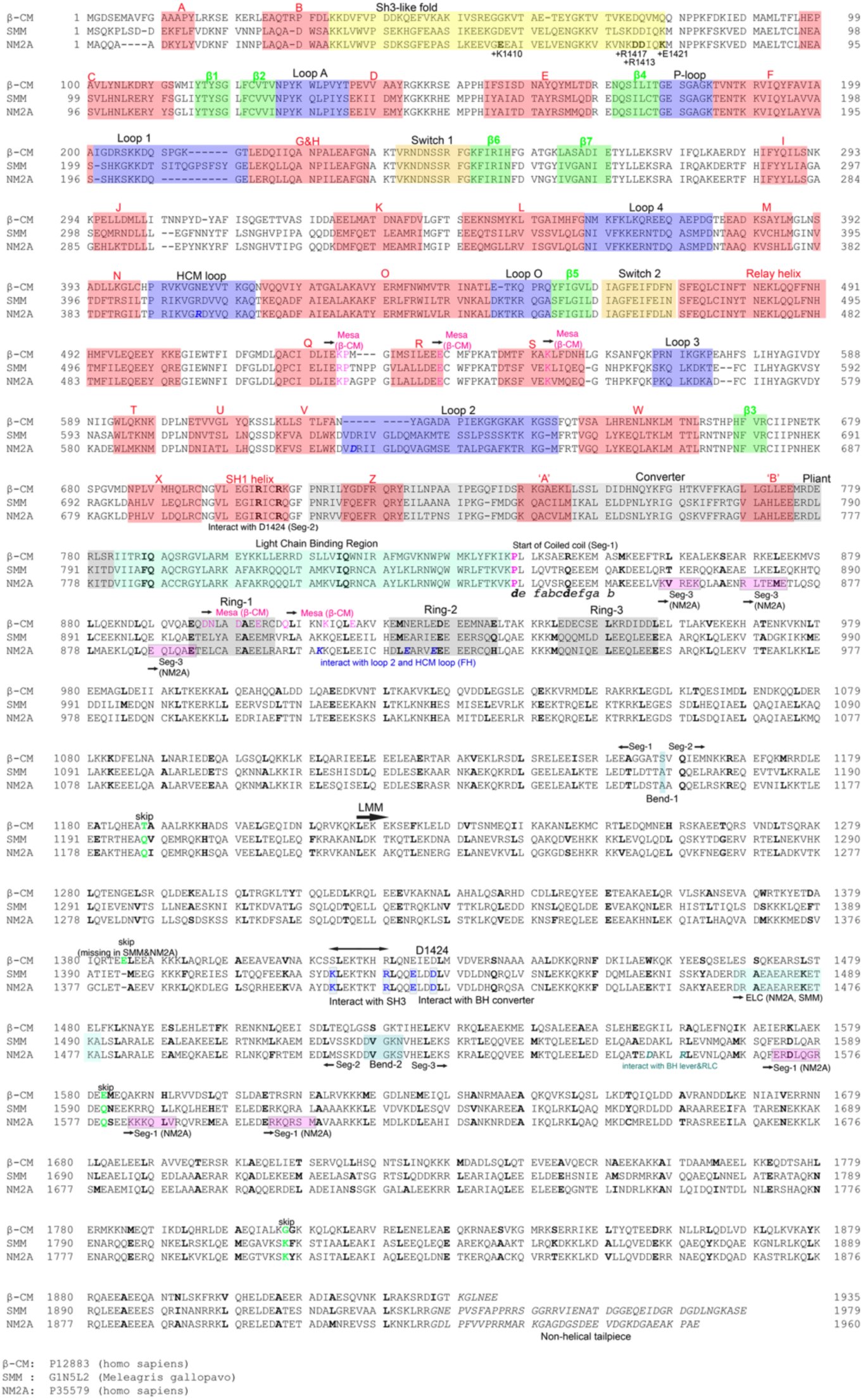
Alignment of β-CM, SMM and NM2A heavy chains. Sequences used are shown below the alignment. Regions within the motor domain, the light chain binding region and the coiled coil are annotated as shown. Bold residues from the invariant proline (magenta) are the ‘*d*’ residues of the coiled-coil heptad repeats. Positions of bends and ‘skip’ residues as indicated. Magenta ‘Mesa’ residues, indicate interacting residues in the mesa trail in β-CM HMM. Magenta boxed regions: interacting regions between Segs-1 and Seg-3 as shown in Fig. 3 for NM2A. Additional sites of interaction as indicated.

**Fig. S7:**
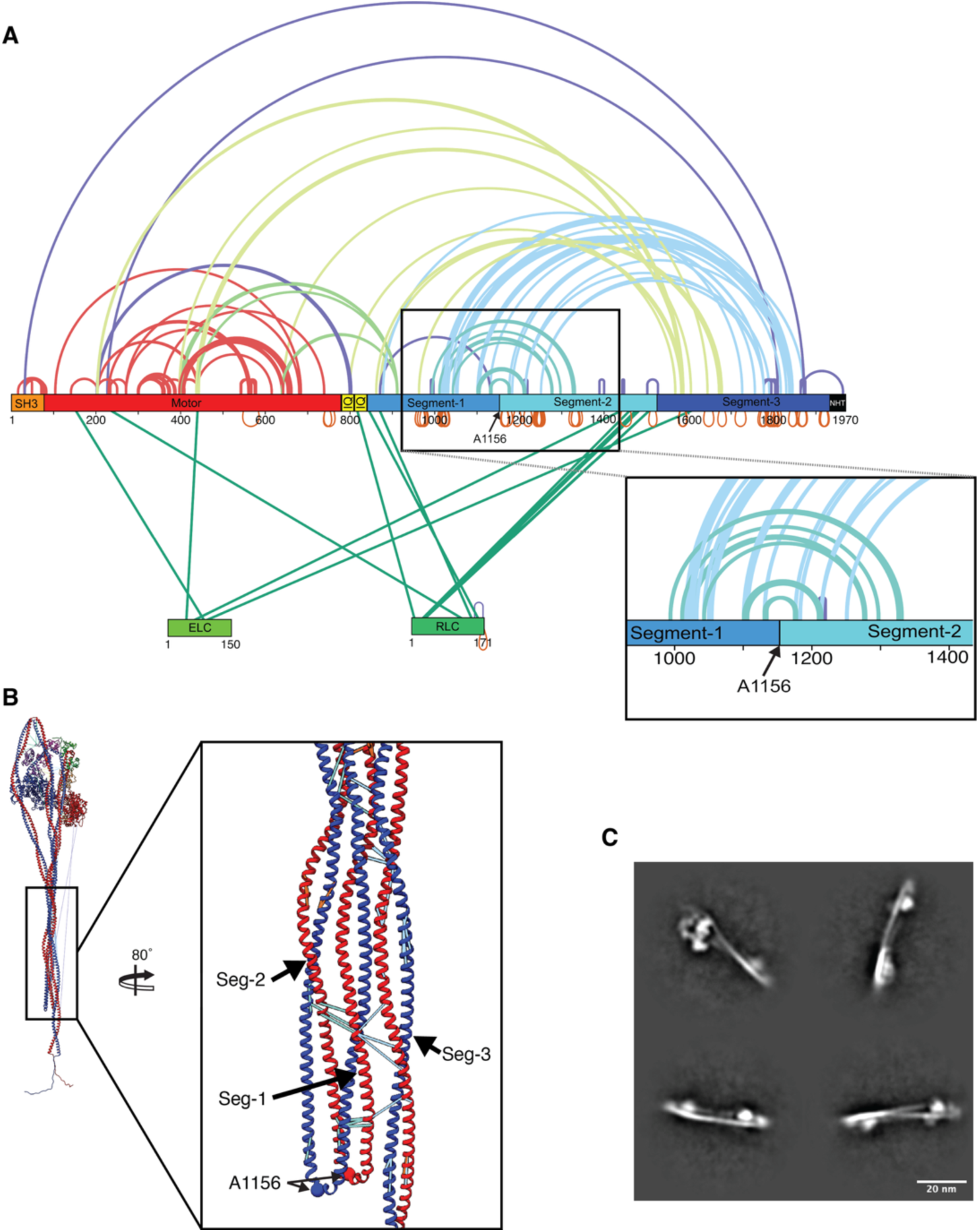
XLMS data for shutdown NM2A. (**A**) Linear diagram to show the domains of NM2A (SH3-like fold, motor etc). Crosslinked residues are indicated by the colored lines drawn. Colors show interactions within domains: red lines for cross-links within the motor, light blue lines between Seg-1 and Seg-3 of the coiled coil, blue-green between Seg-1 and Seg-2 of the coiled coil, light green lines between regions of the motor or coiled coil with light chains, light green for cross-links between motor and coiled coil, and purple for other cross-linked peptides. Orange cross-links signify self-linked peptides. (**B**) shows the shortest cross-link chain pair (under 27Å cut-off; colored according to A) on the shutdown molecule for the boxed region to show the X-links between Seg-1, -2 and -3, placed on the full-length model structure for this region of the structure. **C**. 2D class averages of anti-parallel NM2A dimers (which encompass ∼1% of the cryoEM dataset) that explain very long cross-links outside of the 27Å cut-off within the XLMS dataset.

**Fig. S8:**
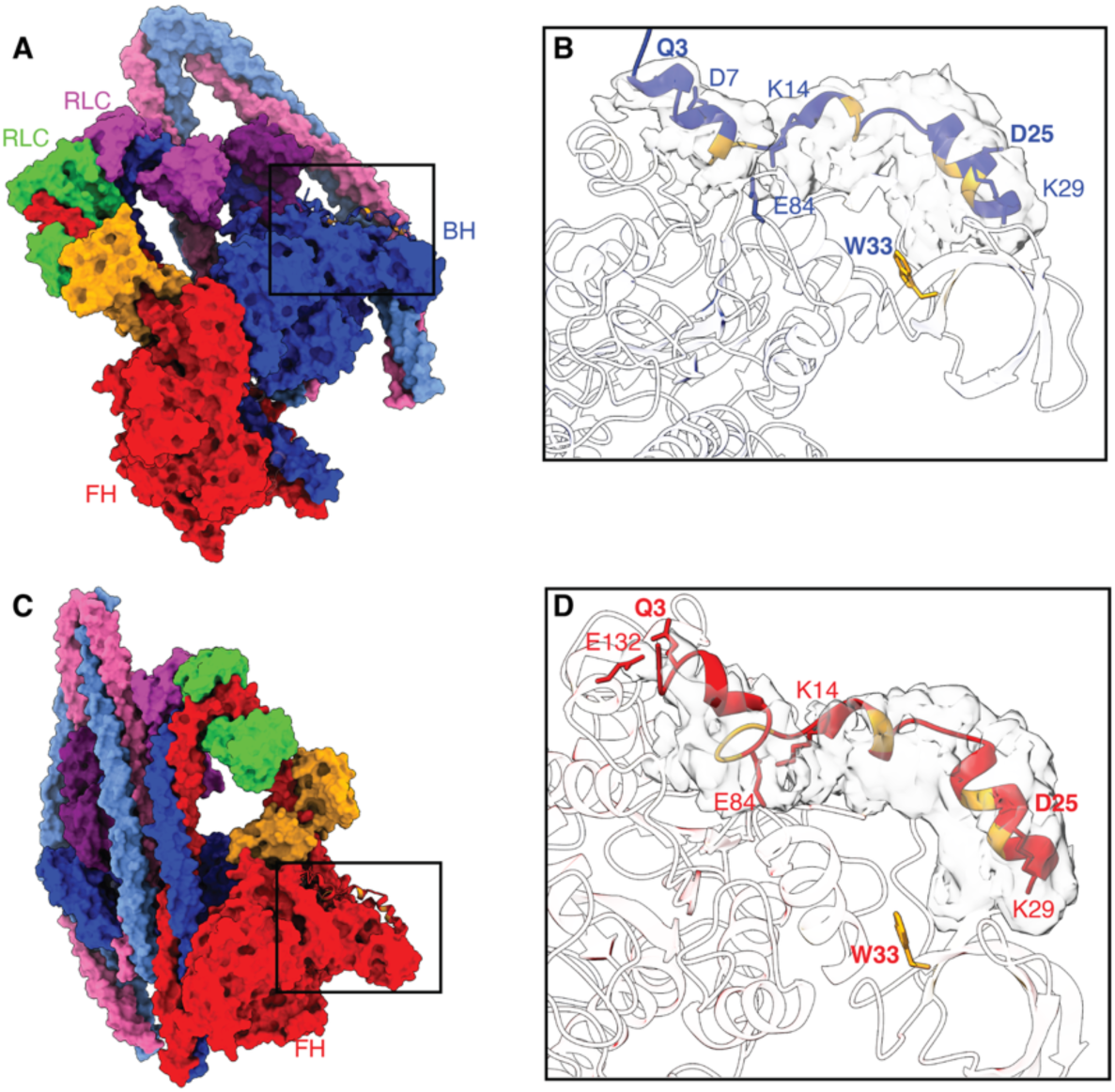
N-terminal domains for Blocked Head and Free Head. (**A**) Colored surface view of the heads region to show the location of N-terminal domain of the BH heavy chain (**B**) Zoomed view of region boxed in **A**. Side chains of interacting residues are shown, with hydrophobic residues in this interface colored in orange. Q3, D25 and W33 are known sites of disease-causing mutations and are labelled in bold text. (**C, D**) Equivalent views as **A, B** for the N-terminal domain of the FH.

**Fig. S9:**
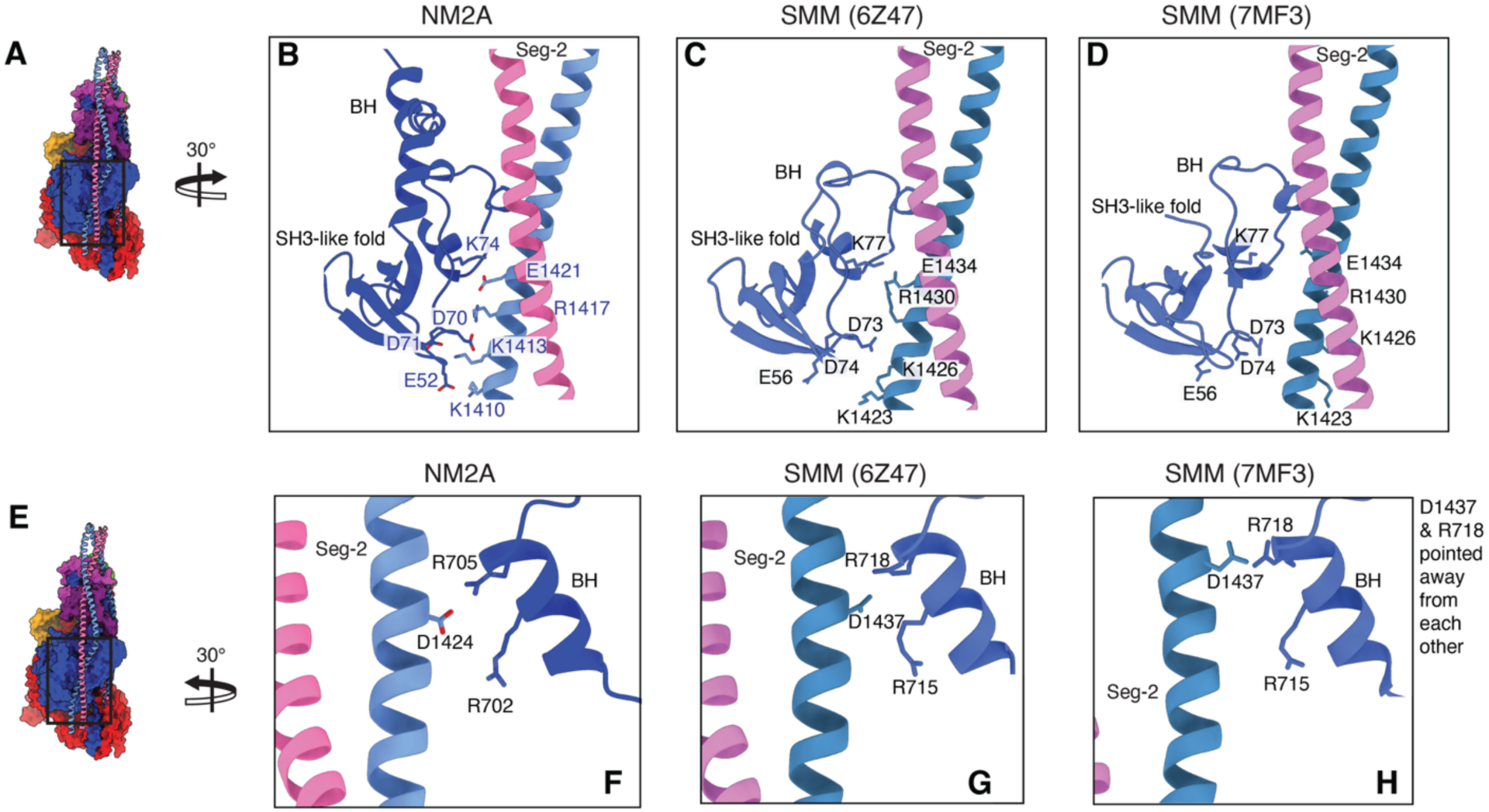
Comparison of Seg2-SH3 and Seg2-SH1-helix interactions in NM2A with SMM. (A) Colored surface view of the heads region (side view) to show the location of Seg-2 interacting with the groove formed within the BH heavy chain (B) Rotated and zoomed view of region boxed in A. Side chains of interacting residues between the SH3-like fold and Seg-2 in the BH are shown. (C, D). Same representation as in B for SMM (PDB: 6Z47) and SMM (PDB:7MF3) structures (aligned with NM2A). (F) Rotated and zoomed view of region boxed in E for NM2A. Side chains of interacting residues between the SH1-helix and Seg-2 in the BH are shown. All three residues shown are known to have disease causing mutations. (G, H). Same representation as in F for 6Z47 and 7MF3 SMM structures.

**Fig. S10.**
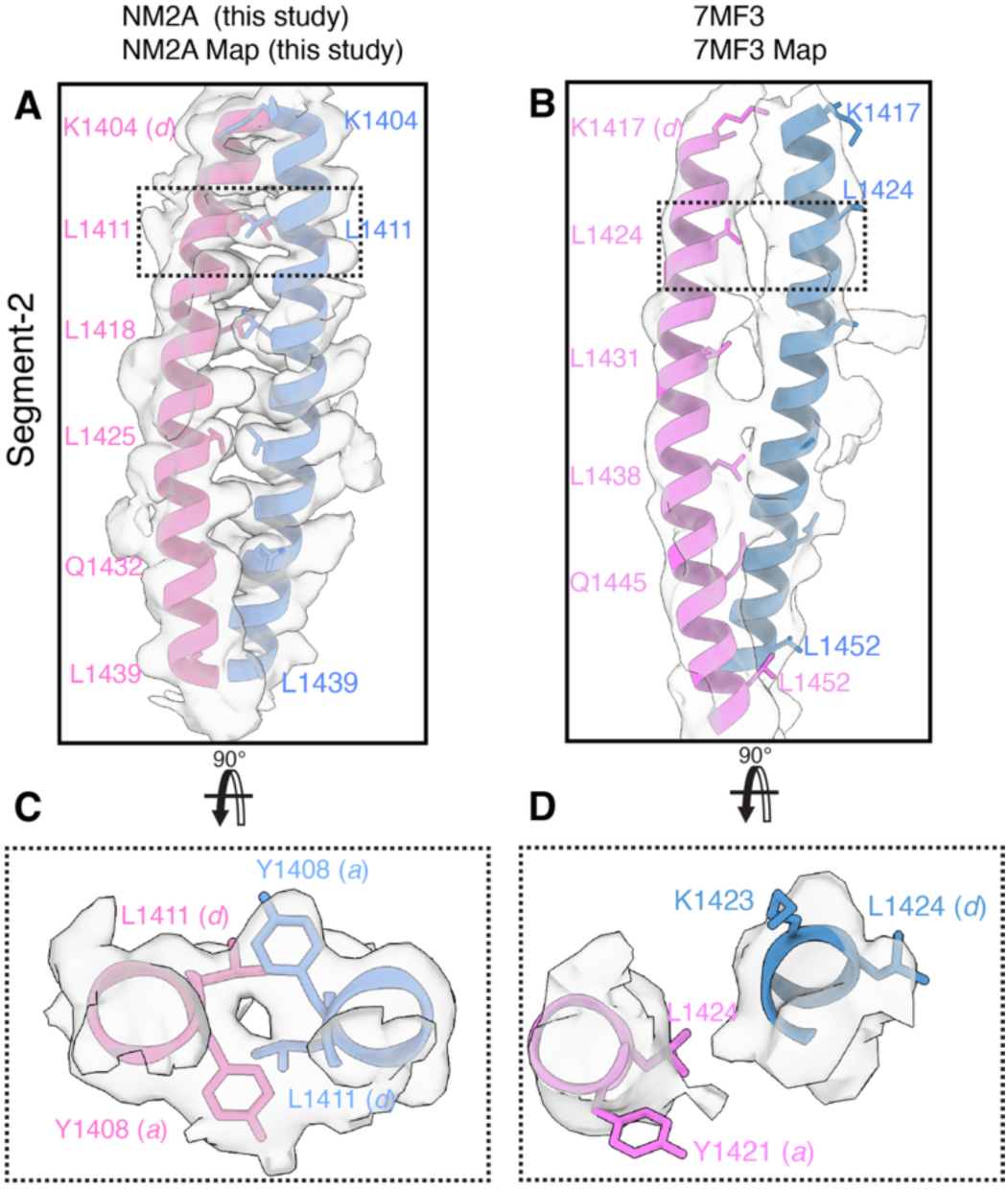
Comparison of the coiled-coil models in Seg-2 of NM2A and SMM (7MF3) **(A)** Representative 5 heptads of coiled coil in Seg-2 (chains G/H:1404-1439) of NM2A in the corresponding segmented cryo-EM map (zoned density displayed at a level of 0.02) with side chains of *d* residues of the hydrophobic seam shown. (**B**) Equivalent representation of the corresponding section of Seg-2 in SMM (PDB:7MF3) in its respective map (EMD-23810, zoned density displayed at a level of 0.1). (**C**) Boxed region from (**A**) to show residues within the hydrophobic seam within corresponding density map (zoned density displayed at a level of 0.1). The *a-a* and *d*-*d* pairing in the hydrophobic seam. (**D**) Boxed region from B to show residues within the hydrophobic seam within corresponding density map (zoned density displayed at a level of 0.1) showing aberrant positions of *a* and *d* residues.

**Fig. S11.**
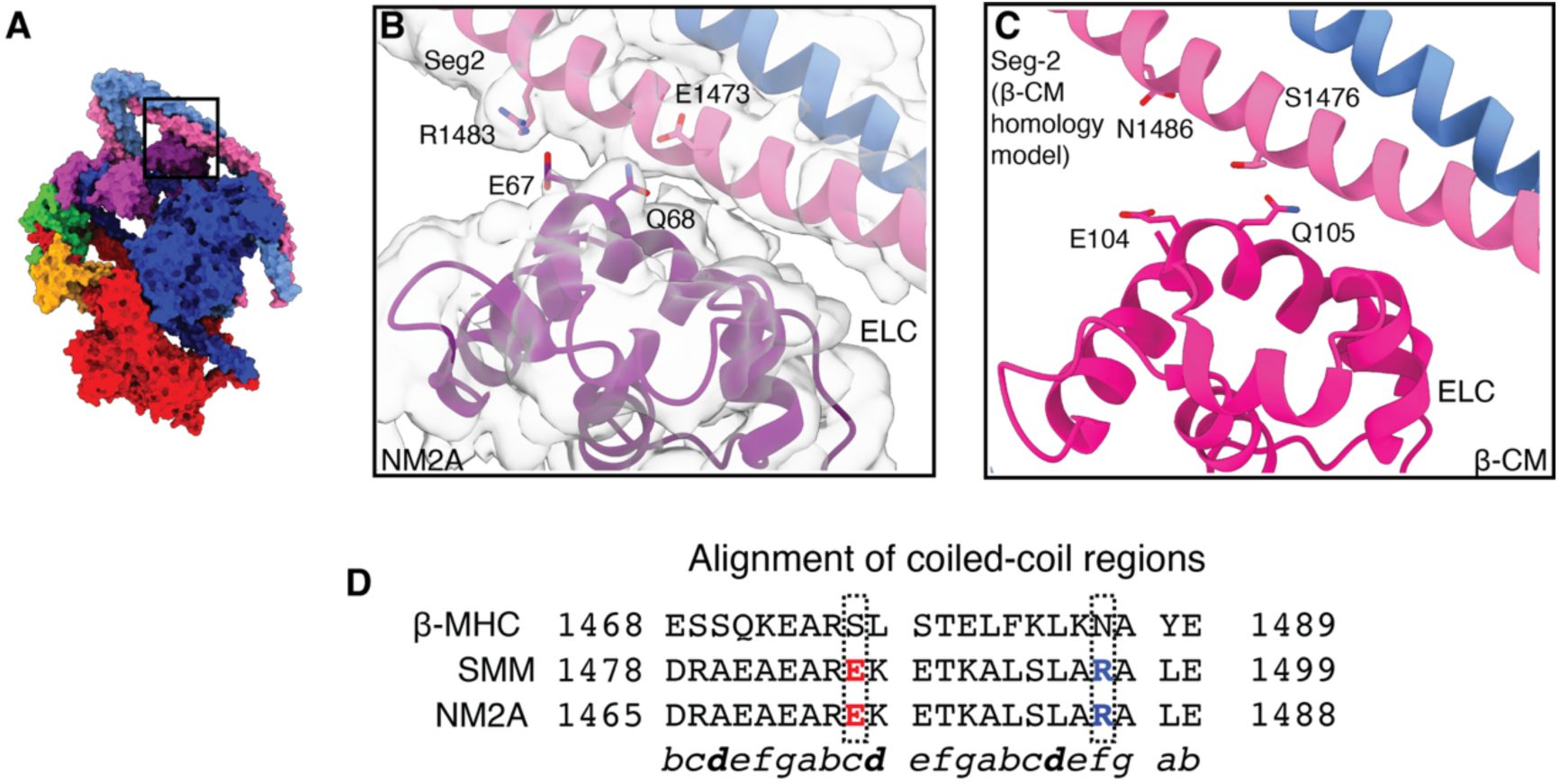
Blocked Head ELC Seg-2 interactions. (A) Colored surface view of the heads region to show the location of BH-ELC Seg-2 interface (B) Zoomed view of region boxed in A. Side chains of interacting residues are shown within the segmented map (C) Equivalent representation of β-CM BH-ELC and a homology model for Seg-2 of β-CM, with corresponding β-CM BH-ELC residues shown (D) Sequence alignment for human heavy chain sequences for β-CM, SMM, and NM2A for the interacting portion of Seg-2. The interacting residues are indicated in bold and color. The heptad position is shown below the alignment. Interacting residues in Seg-2 are not conserved in β-CM.

**Fig. S12:**
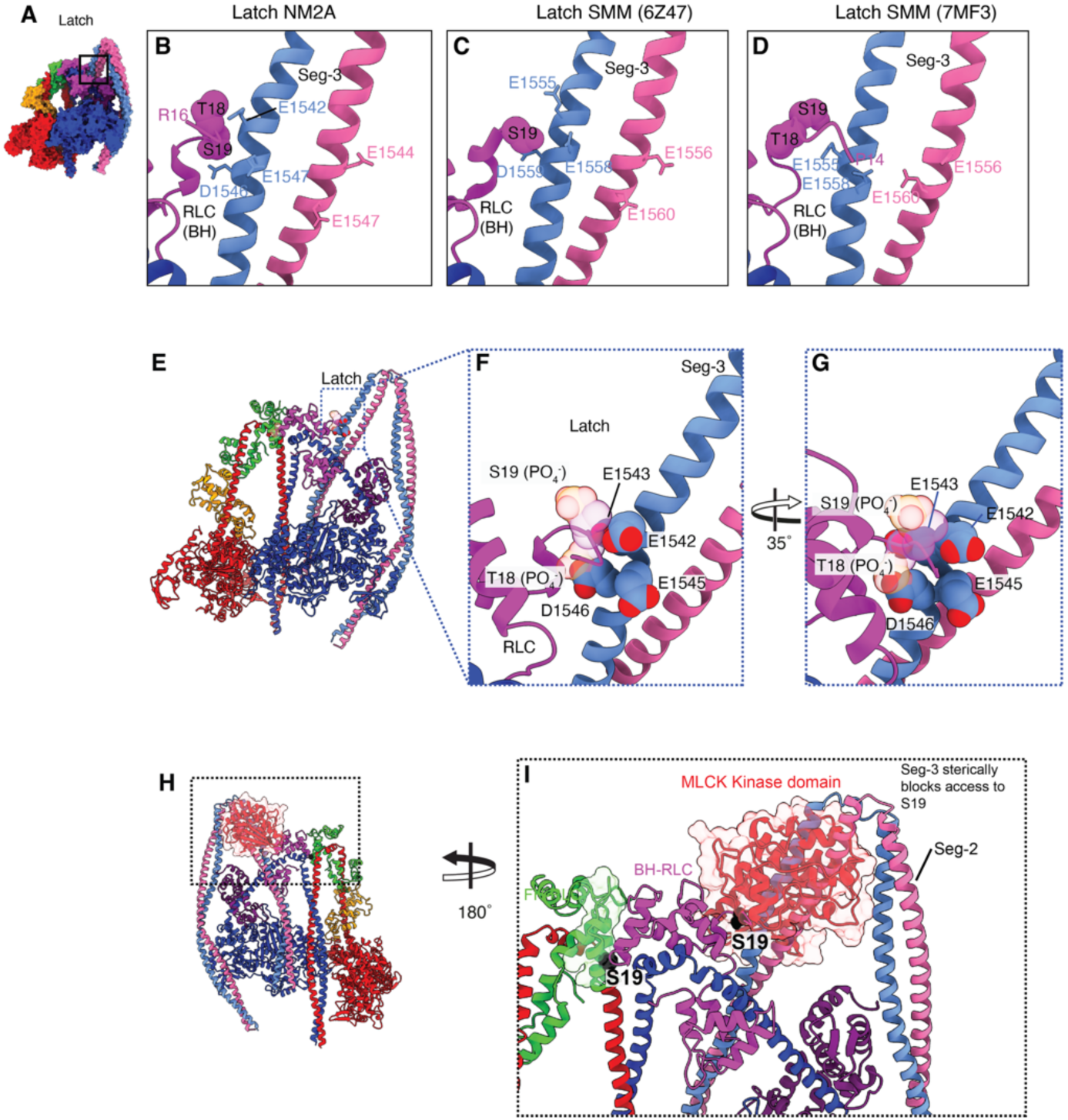
Comparison of latch structures, effect of phosphorylation and MLCK kinase domain interaction. **(A)** Colored surface view of the heads region (back view) to show the location of the BH RLC N-terminal extension (Latch). **(B)** Zoomed view of region boxed in **A.** Side chains of interacting residues between the BH-RLC and Seg-3 are shown. Side chains of the phosphorylatable S19 and T18 residues are shown as spheres. **(C, D).** Equivalent views as in **B** for 6Z47 and 7MF3. The side chains of negatively charged residues of Seg-3 for SMM are shown as sticks. **(E)** Ribbon structure of the modelled phosphorylated molecule showing the position of the latch (boxed region) (**F, G)** Zoomed in views of the phosphorylated latch. Phospho-S19 and Phospho-T18 residues are shown as spheres. Negatively charged residues in the local environment are shown. (**H**) BH RLC N-terminal extension (Latch) docked with the kinase domain of MLCK with the RLC of the kinase-RLC AlphaFold model aligned as in Fig. S3. Boxed region in (**I**), shows how the kinase is sterically occluded by Seg-3 on the blocked-head and unable to interact with S19 in the latch.

**Table S1.**
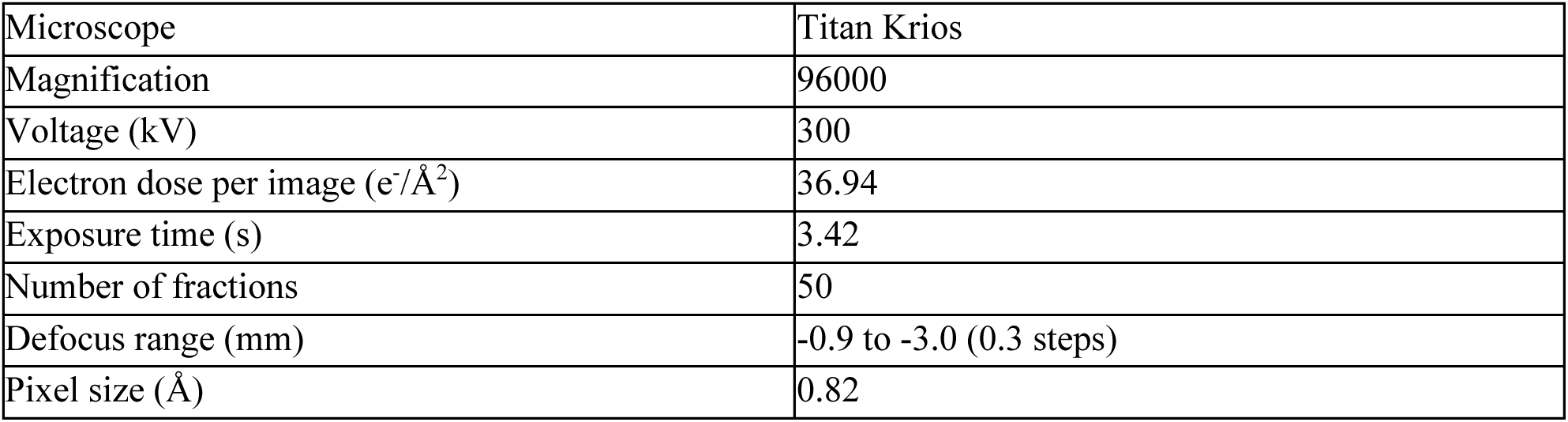
Microscope parameters. Cryo-EM data collection statistics.

### Supplemental Movies

**Movie S1. Shutdown state of non-muscle myosin 2a** (A) Cryo-EM density of NM2A (grey) shutdown heads region at an average resolution of 3 Å. Colored map representation showing heavy chain (blue, red), ELC (purple, orange), and RLC (magenta, green) in complex with segments 2 and 3 (light blue, pink) and fitted pseudo-atomic model. (B) Closeup view of the head tail junction. (C) Close up view of the free-head RLC mortar (dark green) with potentially interacting residues (sticks) in both RLCs shown. (D) MLCK kinase domain docked onto the S19 of the FH RLC showing few steric clashes with the free head.

**Movie S2. Three principal components from 3D variability analysis of NM2A.**

The movie shows conformational heterogeneity of NM2A revealed by 3D variability analysis in CryoSPARC. Principal components 1–3 (PC1–PC3) capture the dominant continuous motions within the dataset. Each component is displayed sequentially in series between the two extreme conformations along that component, highlighting the major structural changes within the molecule.

**Movie S3**. **Tail Flexibility.** The movie shows two-dimensional class averages of non-muscle myosin-2A (NM2A) molecules, arranged to illustrate the range of conformations reflecting in-plane flexing of the tail region. Scale bar, 20 nm.

